# GPCRchimeraDB: A database of chimeric G protein-coupled receptors (GPCRs) to assist their design

**DOI:** 10.1101/2024.12.16.628733

**Authors:** Charlotte Crauwels, Adrián Díaz, Wim Vranken

## Abstract

G protein-coupled receptors (GPCRs) are membrane proteins crucial to numerous diseases, yet many remain poorly characterized and untargeted by drugs. Chimeric GPCRs have emerged as valuable tools for elucidating GPCR function by facilitating the identification of signaling pathways, resolving structures, and discovering novel ligands of poorly understood GPCRs. Such chimeric GPCRs are obtained by merging a well- and less-well-characterized GPCR at the intracellular limits of their transmembrane regions or intracellular loops, leveraging knowledge transfer from the well-characterized GPCR. However, despite the 212 chimeric GPCRs engineered to date, the design process remains largely trial-and-error and lacks a standardized approach. To address this gap, we introduce GPCRchimeraDB (https://www.bio2byte.be/gpcrchimeradb/), the first comprehensive database dedicated to chimeric GPCRs. It catalogs 212 chimeric receptors, identified through literature review, and includes 1,755 class A natural GPCRs, enabling connections between chimeras and their parent receptors while facilitating the exploration of novel parent combinations. Both chimeric and natural GPCR entries are extensively described at the sequence, structural, and biophysical level through a range of visualization tools, with annotations from resources like UniProt and GPCRdb and predictions from AlphaFold2 and b2btools. Additionally, GPCRchimeraDB offers a GPCR sequence aligner and a feature comparator to investigate differences between natural and chimeric receptors. It also provides design guidelines to support rational chimera engineering. GPCRchimeraDB is therefore a resource to facilitate and optimize the design of new chimeras, so helping to gain insights into poorly characterized receptors and contributing to advances in GPCR therapeutic development.

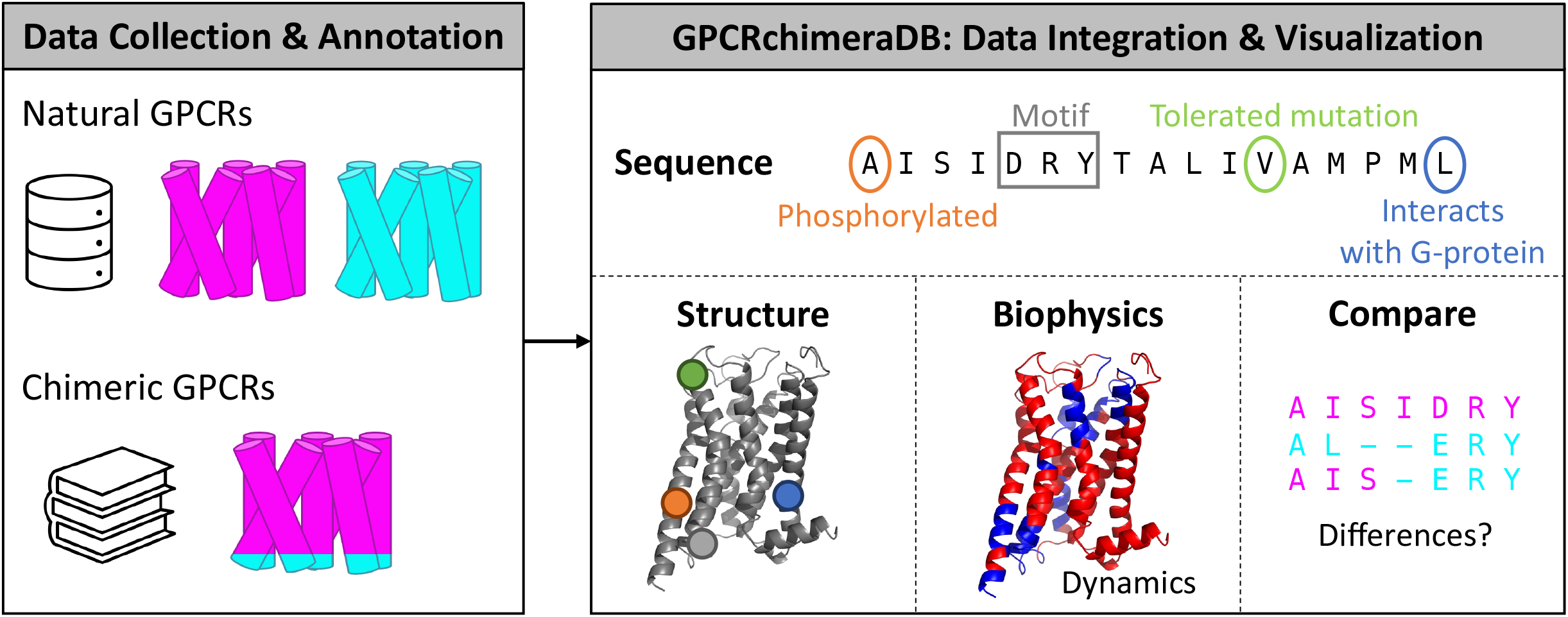

## Introduction

G protein-coupled receptors (GPCRs) are the largest superfamily of membrane proteins. With over 800 GPCRs identified in humans, these receptors make up more than ~4% of the human genome^1,2^. GPCRs serve as signal transducers, allowing cells to respond to a variety of external stimuli, including hormones, neurotransmitters, and sensory signals^1,2^. Consequently, mutations that affect their function can lead to a wide range of diseases, from neurodegenerative and psychiatric disorders to cardiovascular conditions and cancers^3,4^. However, despite their importance, 60 to 85% of the therapeutically relevant GPCRs remain undrugged^5^. Much of the difficulty in targeting GPCRs arises from their innate conformational flexibility, undefined endogenous ligands (orphan GPCRs), insolubility, cell-type specific expression, biased signaling and off-target effects of their ligands^1,6,7^.

Chimeric GPCRs (or hybrid GPCRs) are powerful tools for advancing our understanding of GPCRs^8–10^. Their goal is to leverage what is known about one GPCR to gain deeper insights into the other, less understood, GPCR. They so provide insights into the roles of specific residues in GPCR function^8,9,11^, help decipher biological pathways^8,11,12^, facilitate the elucidation of 3D structures in distinct activation states^7,13^, and assist in identifying novel ligand binders^7^. Chimeric GPCRs are designed by merging regions from 2 natural GPCRs, typically combining a well- and less-well-characterized GPCR. The cutting points, or junctions where the parent GPCRs are fused, are usually positioned at the intracellular (IC) limits of their transmembrane (TM) regions or at their IC loops^9^ (Figure 1A-B). As such, the GPCR on the extracellular (EC) side, the EC parent, provides the EC loops and most of the TM regions while the GPCR on the IC side, the IC parent, supplies the IC loops and possibly, some of the TM regions. Depending on the design strategy, the EC parent may also contribute one or more IC loops, meaning that not all IC loops necessarily originate from the IC parent. However, the IC parent never contributes to the EC side.

**Figure 1:**
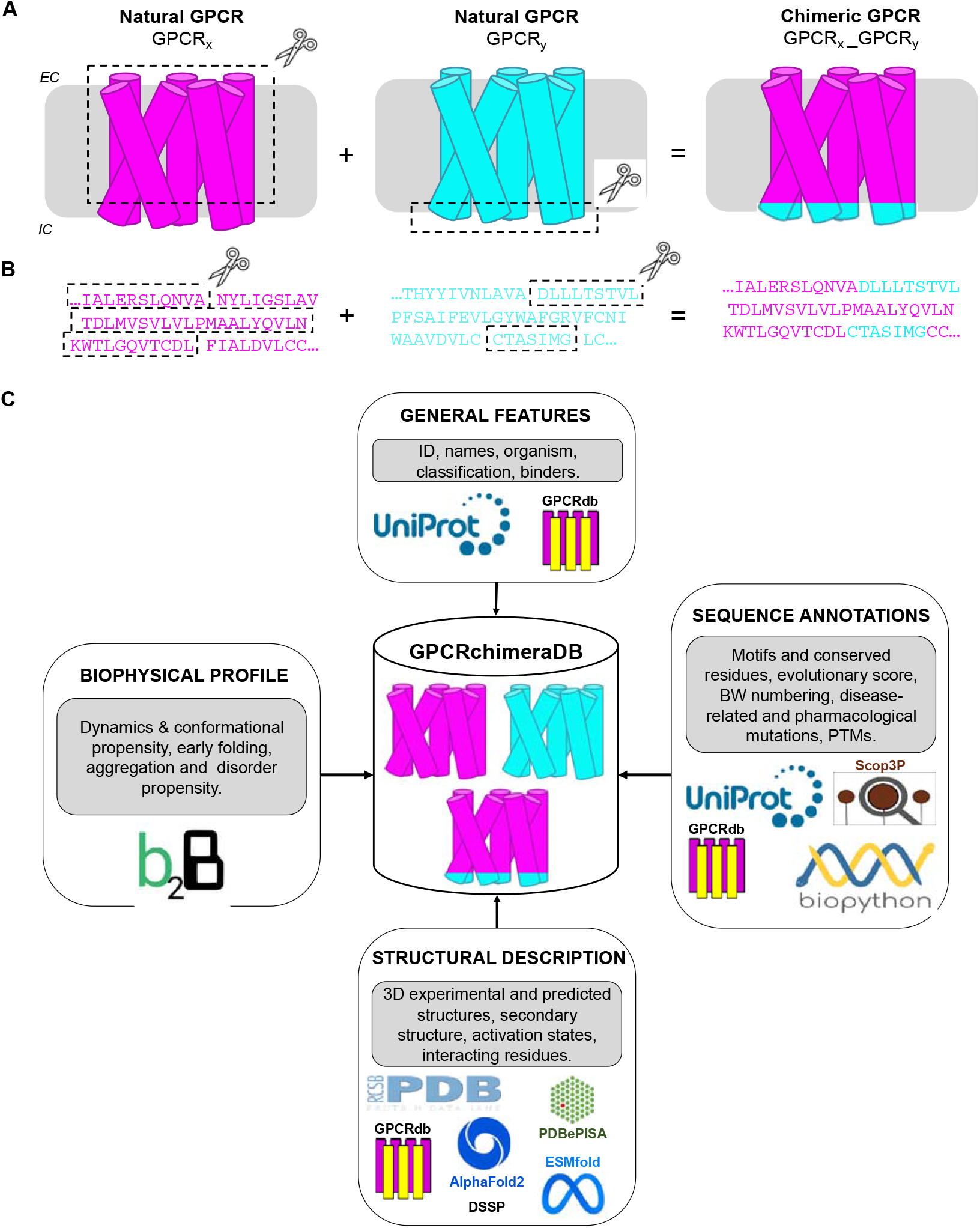
GPCRchimeraDB framework. A. Schematic illustration of chimeric GPCR design. A chimeric GPCR is obtained by merging regions from 2 natural GPCRs, the parents. The EC parent is colored in magenta while the IC parent is colored in cyan. B. Introduction of the cutting points or sites, in the parent GPCRs to obtain the chimeric sequence. The cutting points are typically located at the intracellular limits of the TMs or IC loops. C. Tools and databases used to collect and obtain the annotations available in GPCRchimeraDB to describe the natural and chimeric entries. The data was either extracted from a pre-existing database or generated with a predictor.

Chimeric GPCRs capitalize on the common structure-function relationship among GPCRs^10,11,14,15^. Despite sharing overall less than 30% sequence identity, GPCRs exhibit a highly conserved 3D core structure consisting of 7 TM helices that span the cell membrane, connected by 3 EC and IC loops^1,2^. The key to successful chimeric design lies in selecting optimal cutting sites within the parent proteins to preserve the functional integrity of the EC+TM and IC fragments^8,11,14^: the binding of an agonist at the EC side needs to trigger conformational changes in the TMs to enable the binding of a G protein or β-arrestin at the IC side and allow signal transmission. This requires careful consideration of the sequence, structural, and biophysical (e.g. dynamic) features of the parent receptors as demonstrated by Tichy et al.^11^, the only known study where chimeras were rationally designed. To date, over 212 engineered chimeric GPCRs are available in the public domain. However, in spite of their relevance, no generic design approach to create chimeric GPCRs has been established yet, perhaps due to the lack of overview and annotations in the previous designs.

Here we present GPCRchimeraDB (https://www.bio2byte.be/gpcrchimeradb/), a comprehensive database designed to describe and visualize previously constructed chimeric GPCRs, so facilitating the creation of novel ones. As the first resource to standardize and consolidate data on chimeric GPCRs, it features 212 curated entries derived from extensive literature review. Additionally, it includes 1,755 natural class A mammalian GPCRs, enabling links between chimeric designs and their parent receptors. By integrating detailed annotations, visualization tools, and predictive insights for both chimeric and natural GPCRs at the sequence, structural and biophysical level, GPCRchimeraDB builds upon and extends the descriptions provided by GPCRdb^16^. Moreover, it features tools to compare and analyze chimeric and natural GPCRs as well as guidelines to support the design of novel chimeras. GPCRchimeraDB opens new opportunities for researchers to explore the field of GPCR chimeras and consider their potential as tools to investigate poorly characterized GPCRs of interest.

## Material and methods

### Collecting natural and chimeric GPCRs

The natural GPCR entries in GPCRchimeraDB consist of all reviewed class A (rhodopsin-like) mammalian GPCRs available on InterPro^17^ (IPR017452). Based on models available from the AlphaFold Protein Structure Database^18^ (AFDB), entries with more or fewer than 7 predicted TMs were removed.

Chimeric GPCR entries were retrieved from the literature using a systematic search on databases like PubMed and Google Scholar. The search included keywords such as ‘GPCR chimera’, ‘GPCR hybrid’, ‘GPCR design’, and ‘GPCR engineering’. Cross referencing was also critical to retrieve previously designed chimeras. In addition, a BLAST^19^ search (E-value 0.01) against the Protein Data Bank^20^ (PDB) was performed to identify chimeric GPCR chains. Human ADRB2, ADRB1, ACM2, OPRM, OPRK, AA2AR, and AGTR1 were used as query because conformation-stabilizing VHHs (Nanobodies)^7^ have been designed for these GPCRs and it is therefore more likely that such GPCR is involved in chimeras (see ‘Type 3 chimeras’ in ‘Classifying chimeras in GPCRchimeraDB’).

For each identified chimeric design, the following information was extracted from the related paper:

- Protein sequence: When the DNA sequence was provided, it was translated into a protein sequence using Python scripts^21^.
- UniProt^22^ IDs of the 2 parent GPCRs: If not provided, a BLAST search against the SwissProt database (E-value 0.001) was performed on the chimeric sequence to identify the parent GPCRs. Based on the section the parent GPCRs provided, we defined the EC and IC parent (see definition in the Introduction).
- Positions of the cutting sites: If not described, a pairwise local sequence alignment using Biopython^23^ was performed between the chimeric and the parent sequences (using the ‘pairwise2.align.localxx’ function).
- Mutations introduced into the chimeric sequence compared to the parent sequences.
- Experimental 3D structure(s) of the chimera (PDB ID), if available.
- Functionality information: Functionality data is included as either a quantitative value (when available) or a qualitative value (yes/no). This data is not normalized across the entries (see section ‘Previously designed chimeras in GPCRchimeraDB’).
- Binding ligand(s) and G protein(s), if available.
- Design type (Type 1/2/3 as defined in ‘Classifying chimeras in GPCRchimeraDB’) and design purpose of the chimera (application).
- DOI of the paper.

Duplicate designs are considered as the same entry in the database but are referred to by multiple DOIs. This data retrieval and formatting was handled by an in-house Python pipeline.

### Description of the entries

For every natural and chimeric entry, a comprehensive set of features was collected and formatted using a Python pipeline divided into 4 sections: the general features (e.g., classification and ligands), the sequence annotations, the biophysical profile, and the structural description. These sections are described in more detail below, with the features and tools summarized in Figure 1C. The retrieved data was compiled into uniform JSON files, one per entry (see Zenodo for an example). These files were then uploaded individually to GPCRchimeraDB. This system simplifies updating entries when new information becomes available (more details in the section ‘Technical architecture’).

#### General features

For every natural GPCR, we gathered their UniProt ID, protein name (also called gene or abbreviated name), alternative name(s) and source organism (from UniProt) as well as their IUPHAR name (from GPCRdb). Moreover, based on information from GPCRdb, all natural entries were classified into families and subfamilies based on the endogenous ligands they bind. A phylogenetic-based subfamily classification was also added, where we extended the original version by Cvicek et al.^24^ for a limited dataset to all mammals using first CDhit clustering^25^ (40% sequence identity cut-off) and then an annotated phylogenetic tree generated with IQtree^26^. Additionally, information on the principal endogenous ligand(s) and, G protein and β-arrestin subclass(es) the GPCRs bind was also extracted from GPCRdb and added to the database.

Currently, there is no established naming convention or classification system (including class, family, and ligand- and phylogenetic-based subclass) for chimeric GPCRs. To address this in GPCRchimeraDB, all chimeras were given a unique identifier and abbreviated name: <*UniProt ID EC parent*>_<*UniProt ID IC parent* >_i and < *UniProt Protein Name EC parent*>_<*UniProt Protein Name IC parent* >_*i*, respectively and where *i* denotes the order in which we annotated the chimeras (e.g, P43140_P18841_1 and ADA1A_RAT_ADA1B_MESAU_1, respectively). To further describe the chimeric entries, we concatenated the features of their EC and IC parents, in that order. For example, for P02699_P35348_1, this results in Rhodopsin *α*1A-adrenoceptor (name), Bovine & Human (organism), A & A (class) and Opsins & Adrenoceptors (family).

#### Sequence annotations

Class A GPCRs are characterized by a set of well-conserved motifs, also known as microswitches, that are key for the receptor’s function^1^. These motifs include the ionic lock switch (E/DRY/W), the transmission toggle switch (CWxP), the Tyr toggle switch (NPxxY), the PIF motif, and the hydrophobic lock. The presence of these motifs is annotated in GPCRchimeraDB. To support residue-level analysis, Ballesteros-Weinstein (BW)^27^ and GPCRdb numbering schemes^28^ are provided for natural GPCR entries. For chimeric entries, we define their numbering using BW’s methodology: the most conserved residue in each TM (N, D, R, W, P, P, and P in TM1 to TM7, respectively) is designated as x.50, where *x* corresponds to the TM number. The adjacent residues are numbered accordingly, with the preceding residue labeled x.49 and the following residue labelled x.51. These standardized numbering systems enable more informative residue identification and facilitate cross-receptor and -database comparisons. For natural sequences, each residue is also assigned an evolutionary conservation score, reflecting its level of conservation compared to the consensus sequence of its phylogenetic subclass. Additionally, we computed an overall conservation score for each phylogenetic subclass, using Shannon entropy to quantify sequence variability. For chimeras, their sequence is also directly compared to their parents’ sequence on their entry page.

To extend the description of the sequence space of the natural entries, we included mutagenesis information. Disease-related mutations were collected from UniProt together with their predicted functional effects. Pharmacological mutations that are known to alter the response of a GPCR to specific ligand were retrieved from GPCRdb. Finally, post-translational modifications (PTMs) from UniProt and Scop3P^29^ were added as well as disulfide bonds (UniProt).

#### Biophysical profile

The predicted biophysical profile includes the protein’s backbone and sidechain dynamics as well as the early folding, disorder and beta-sheet aggregation propensity and the conformational propensities (helix, sheet, coil and polyproline II helix). We obtained these features using sequence-based predictors that are part of the b2btools^30^. For the chimera entries, we also computed the difference between their biophysical profile and the one of their parents to enable comparison. To achieve this, we aligned the chimera’s and parents’ sequence and calculated the difference between the predicted values for each.

#### Structural description

Based on the PDB ID(s) associated with each UniProt ID, all available 3D experimental structures of natural GPCRs were retrieved from the PDB, along with the method used to obtain the structure and its resolution. The PDB information of chimeric designs with a resolved 3D structure was also retrieved. The GPCR chain was identified based on its annotation on the PDB, and a structure was retained only when its GPCR chain was longer than 200 residues (GPCRs are typically around 400 residues long). When available, the activation state (active, intermediate, inactive) of the 3D structure was also retrieved from GPCRdb. Furthermore, using PDBePISA^31^, we identified in the full 3D structure complex, the GPCR residues interacting with a ligand or an effector (G protein, nanobody/antibody, or β-arrestin). Sequence-to-structural position mapping was performed using the information provided by the PDBe^32^ (mmCIF files). The experimental structural information was further enriched with predicted models from AlphaFold2^33^ (AF2), AlphaFold2 multistate^34^ (AFms) for both natural and chimeric GPCRs, as well as ESMFold^35^ (ESMF) models exclusively for chimeras. These models were either retrieved from existing sources or predicted in-house (see Table S1 for details). Finally, the limits of the TM regions were assigned using DSSP^36^, based on the available experimental 3D structure or, when unavailable, the AF2 models (see ‘Procedure to define the TM limits’ in the Supplementary Material for more details).

### Sequence alignment tool

The sequence alignment tool in GPCRchimeraDB is based on a profile Hidden Markov Model (pHMM). The pHMM is generated on-demand with HMMER^37^, based on a pre-computed Master Alignment (MA) created using the structure-informed MSA pipeline SIMSApiper^38^. The use of a MSA slightly different from the one proposed by GPCRdb is discussed in the Supplementary Material. More details about the alignment procedure and the MA are also available in the Supplementary Material.

### Accessibility and maintenance

GPCRchimeraDB is a freely accessible database under the FAIR principles^39^. The database is currently maintained and will be updated on a regular basis. The database is supported by extensive documentation (https://gpcrchimeradb-docs.readthedocs.io/) and a FAQ (https://www.bio2byte.be/gpcrchimeradb/faqs).

### Technical architecture

The GPCRchimeraDB is deployed inside a GNU/Linux web server running Ubuntu 22.04. The backend component written in Python3 is based on the Django framework to build the endpoints and the database middleware. Internally, this component relies on a SQLite3 file-based database. The database layer stores the JSON files as well as the interpretation data in tables. The backend component has a CLI mechanism to import entries allowing maintainers to update and add new entries. The frontend component built on React.js library fetches the data via HTTP requests to the backend in order to render pages and visual components. The frontend codebase is compiled to static HTML, CSS, and JS files, while the backend component runs as an uWSGI socket managed by a supervisor service. Web server’s HTTP requests are handled by a NGINX HTTP proxy instance which redirects GCPRchimeraDB requests to both frontend pages and backend API endpoints.

## Results

### Classifying chimeras in GPCRchimeraDB

In total, we identified 212 chimeric GPCRs in the literature. Upon investigation, these chimeras can be categorized into 3 distinct design types based on their design strategy and applications. As such, in GPCRchimeraDB, each chimera is assigned a design type:

- Type 1 designs, also called Opsin-based or OptoXRs designs, combine a light-sensitive GPCR, often Rhodopsin, at the EC side of the chimera with a poorly characterized GPCR at the IC side. Hence, through light stimulation, the designed chimera is activated, enabling the study of the biological pathways associated with the IC parent, which could not be activated previously due to the absence of a known ligand (orphan GPCR). This design strategy also enables identification of the importance of specific regions in IC parent, for example for G protein selectivity.
- Type 2 designs have a similar configuration as Type 1 chimeras, but the EC parent is a non-sensory GPCR, i.e., the chimera gets activated by a known endogenous or synthetic ligand instead of light. This again enables the study of the (activated) IC side GPCR or its related signaling pathway(s).
- In contrast, Type 3 contains the well characterized GPCR on the IC side of the chimera. For these IC parents, a stabilizing effector (nanobody/VHH) that can lock the GPCR in a specific conformation is known^7^. This effector can so stabilize the chimera in, for example, a disease related conformation and drug screening for this disease state can then be performed to help deorphanize GPCRs. By stabilizing the chimeric GPCR in specific conformations, structure elucidation can also be facilitated which can, in turn, aid structure-based drug design or bring new structural insights.

To summarize, the design strategy of the chimera depends on the research goal: elucidating GPCR signaling pathways or unravelling key regions for the protein function (Type 1 and 2), identifying new potential ligands (Type 3), or supporting 3D structure determination (Type 3). These 3 design types are illustrated in Figure S2.

### Previously designed chimeras in GPCRchimeraDB

As shown in Table 1 and Figure S3A, 94 of the 212 designs in GPCRchimeraDB are rhodopsin-based chimeras (Type 1), largely inspired by the pioneering work of Kim et al. in 2005^15^. In 2018, Morri et al.^12^ engineered a library of 63 OptoXRs, incorporating orphan GPCRs to gain insights into these poorly characterized receptors, making it the largest single-study dataset of GPCR chimeras. The overall extensive use of rhodopsin underscores its utility as a reliable framework in the design of GPCR chimeras. In contrast, only a few Type 3 chimeras have been successfully constructed (15 in total). They all consist of the human κ opioid receptor (KOR) at the IC side, stabilized by a nanobody (Nb6), and combined with a GPCR of interest at the EC side^7^. A commercial patent^40^ has been filed for the development of Type 3 chimeras which highlights their significant therapeutic and economic potential. Type 2 designs are referenced in the largest number of distinct publications (30 papers in total which describe 103 designs with 42 unique parent combinations) and display the broadest applicability, ranging for example from understanding the role of the C terminal in functional desensitization^41^ to exploring G protein selectivity^42^ and comparing the pharmacology of GPCR homologs^43^. Notably, DREADD-based GPCR chimeras^44^ also fall under Type 2 designs. In these chimeras, the EC parent is a DREADD, a GPCR mutated to respond exclusively to a synthetic ligand with minimal biological activity^45^, while the IC parent is the GPCR of interest. This configuration overcomes challenges such as off-target effects and the poor bioavailability of natural ligands, which often hinder studies of natural and some chimeric designs.

**Table 1:** Data integrated into GPCRchimeraDB. On the left, the data types in the database and on the right, features available to describe the entries. ND: Not Determined, NA: Not Applicable.

| Data | Total | Features | Total Natural GPCRs | Total Chimeric GPCRs |
| --- | --- | --- | --- | --- |
| Natural entries | 1,755 | Disease-related mutations | 310,806 | ND |
| Human entries | 712 | Pharmacological mutations | 30,085 | ND |
| Non-human entries | 1,043 | PTMs | 5,851 | ND |
| Unique EC parents | 43 | Disulfide bonds | 1,859 | 215 |
| Unique IC parents | 99 | Experimental 3D structures | 1,184 | 10 |
|  |  | Predicted 3D models | 2,711 | 848 |
| Chimeric entries | 212 | Activation states | 2,311 | 434 |
| Type 1 chimeras | 94 | Interacting residues | 12,522 | 218 |
| Type 2 chimeras | 103 | Endogenous ligands | 949 | NA |
| Type 3 chimeras | 15 | G protein and β-arrestin | 230 | 145 |

While all chimeras reported in the literature are confirmed to express, some exhibit reduced functionality or are entirely non-functional compared to their parents. In GPCRchimeraDB, each chimera’s functionality is described using either the quantitative or qualitative value provided in the publication describing them. However, in case users want to compare functionality values across different chimeras, we strongly advise them to refer to the original publications and compare the experimental conditions. This is because the assays used to assess functionality can vary (e.g., real-time versus endpoint assays), as do experimental conditions (e.g., reagent concentrations) and the way results are presented (relative versus absolute values). For this reason, it is impossible to standardize this information, and we do not report the total number of successful chimeras in GPCRchimeraDB. Nevertheless, it is important to acknowledge the likely publication bias towards well-expressing and functional GPCR chimeras. To address this gap, GPCRchimeraDB (https://bio2byte.be/gpcrchimeradb/submitDesign) encourages researchers to submit unpublished chimeric designs, to provide a more comprehensive view of previous work and facilitate progress in the field.

Note that to be included as a chimera in GPCRchimeraDB, the exchanged region between 2 GPCRs must span at least 3 residues, otherwise we define the construct as a mutant. Additionally, some constructs found in the literature were not included in GPCRchimeraDB:

- Chimeras designed with natural GPCRs not part of class A. Interclass chimeras involving a class A parent GPCR (e.g. class A-F chimeras) were also not considered.
- Chimeric GPCRs with exchanged EC loops and/or exchanged TMs.

While such alternative designs likely also offer valuable insights into GPCRs, GPCRchimeraDB (v1) focuses exclusively on Class A GPCRs, with chimeras constructed by swapping the IC boundaries of their TMs and IC loops. On Zenodo we have collected the alternative designs encountered during database curation that we might consider for inclusion in future versions.

### Extensive description of chimeric and natural GPCRs in GPCRchimeraDB

In total, 212 chimeric and 1,755 natural class A GPCRs were uploaded in GPCRchimeraDB (Figure S3B). For natural GPCRs entries, the UniProt ID allowed the retrieval of experimental and precomputed results from various databases (UniProt, GPCRdb, PDB, AFDB, Scop3P and, precomputed AFms models), which were then further enriched with predicted data (b2btools and DSSP, Figure 1). On the contrary, data for chimeric GPCRs was directly extracted from the original publications describing them. When available, corresponding experimental 3D structures were retrieved from the PDB. Additionally, these entries were supplemented with predicted data (b2btools, DSSP and, AF2, AFms and ESMF models). Information sourced from UniProt, GPCRdb and Scope3P (mutagenesis and PTMs, in particular) is therefore not available for chimeras. As a result, chimeric GPCRs have more limited annotations compared to natural GPCRs (Table 1). Nevertheless, GPCRchimeraDB offers a broad description of both the chimeric and natural entries.

For every entry, a brief overview (classification, binding partners, …) is presented next to an interconnected interactive sequence-structure (1D-3D) viewer (Figure 2A-C). The 1D-3D viewer enables studying the location of pharmacological and disease-related mutations, conserved motifs, interacting residues or PTMs, directly on a user-selected 3D structure. This 3D structure can be either experimentally resolved (from the PDB, when available) or predicted (AF2, AFms and ESMF). Also, when available, the user can also choose the 3D structure based on its activation state (active or inactive, defined using GPCRdb’s definition or by AFms). For each entry, an AF2 model was either retrieved or generated (see Table S1). However, AFms models are currently available only for chimeric and human natural GPCRs. AFms models, particularly in the active state, have demonstrated higher accuracy in TM regions compared to standard AF2 models, as AF2 tends to be biased toward predicting the inactive state^34,46^. This bias likely stems from an unbalanced training set and template database. GPCRdb’s latest update (2025) has partially addressed this issue by recomputing human AF2 and AFms models using the most recent PDB structures. Computing the missing AFms models for non-human natural GPCRs and updating the AF2 non-human models will be considered in future updates of our database.

**Figure 2:**
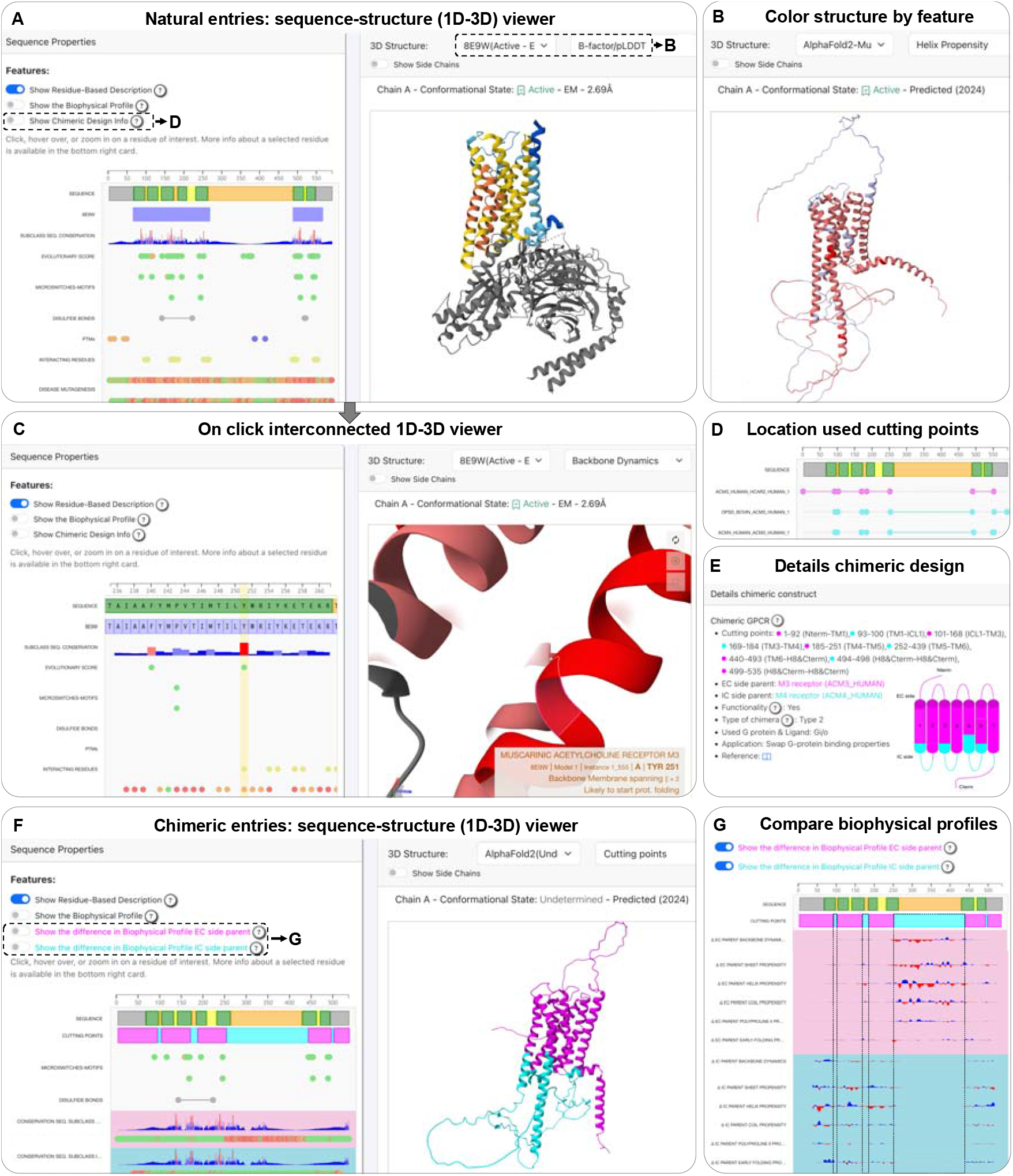
Overview of GPCRchimeraDB’s interactive visualization features. A. Default view of the interconnected sequence-structure viewer on each natural entry page. B. A 3D structure can be selected from a list of experimental and predicted models, with color schemes based on B-factor/pLDDT or biophysical features. C. Clicking a residue in the sequence or structure viewer zooms in for detailed analysis. D. Location of other cutting sites in designed chimeras involving the selected natural entry. E. Detailed description of the chimera on its entry page. F. Default sequence-structure viewer for each chimera entry page. G. Feature to compare the biophysical profile of the chimera with its parents.

For chimeric entries, their ESMF model is also provided. ESMF generates 3D models directly from single sequences, leveraging a language model^35^. In contrast, AF2 requires evolutionary information from a MSA and structural templates^33^. Therefore, the AF2 model of chimeric GPCRs may be biased by the evolutionary and structural information of the EC parent, which contributes more significantly to the chimera’s sequence than the IC parent. In Figure S4, we compared all available GPCR chimeric experimental 3D structures (10 in total, representing 7 Type 3 chimeric designs) to their predicted models (AF2, ESMF, and AFms). However, this analysis does not allow us to determine the most suitable predictor for chimeric GPCRs in general: (i) the results are inconclusive, with relatively small and variable RMSD differences between the predictors and the PDB entries, (ii) all chimeras in this study have only one swapped region (end of TM5–beginning of TM6) but the number and length of swapped regions likely impact prediction accuracy, and (iii) all chimeras share the same IC parent (the *κ* opioid receptor) and only 6 different EC parents, each with experimentally resolved 3D structures in the PDB, which likely influence AF2-based predictions in this benchmark. Given the dataset’s limited size and lack of heterogeneity, we leave it to the user to select the most appropriate model for their specific application. AF2 single-sequence mode (default parameters with and without relaxation on ColabFold^47^) was also tested. While it performs well on *de novo* designs^48^, it produces highly inaccurate models for chimeric GPCRs, likely due to the absence of evolutionary information (Figure S5).

In addition, the biophysical profile of all GPCRs is provided on their entry page. It represents emergent features that evolution conserves to maintain protein function, such as backbone dynamics or conformational propensities, that are not directly apparent from protein sequence or structure (see the ‘Biophysical profile’ section for more details). This type of information can help detect global homology between proteins^49^, as well as improve the pairwise alignment of proteins that are evolutionary distant^50^, and is therefore particularly relevant to study GPCRs as their sequence identity can be low, especially in their loop regions. These biophysical profiles can be directly mapped on a selected 3D structure to facilitate interpretation (Figure 2B) and are unique to GPCRchimeraDB.

For the chimeric entries, the EC and IC parents, design type, functionality value (qualitative or quantitative), application, and the location of the cutting sites are specified on their entry page. A schematic representation of the chimera is also generated (Figure 2E), and the location of the cutting sites can be visualized directly on the 3D structure (Figure 2F). Additionally, their biophysical profile and sequence can be compared to those of their parents (Figure 2F-G) and a table listing other designs with at least 1 parent in common is available. Finally, for the natural entries involved in chimeric designs, all the previously used cutting points are highlighted (Figure 2D) and a table listing all the chimeric designs they are involved in is provided.

### Leveraging GPCRchimeraDB: How to design a novel chimeric GPCR?

In this section, we describe how to utilize GPCRchimeraDB to guide the design of a novel chimera involving GPCRx, the GPCR under investigation. The design process begins by determining the appropriate design type (Type 1, 2 or 3, see ‘Classifying chimeras in GPCRchimeraDB’). This dictates whether GPCRx will serve as the EC or IC parent of the chimera:

- Type 1 or 2: For studying the functionality or signaling pathway of GPCRx. GPCRx will be the IC parent.
- Type 3: For uncovering the 3D structure or conducting ligand screening of GPCRx. GPCRx will be the EC parent.

The next step involves selecting GPCRy, the second parent. GPCRy should be a well-characterized receptor that complements the lack of knowledge for GPCRx. Depending on the design type, it is important to choose a GPCRy for which either a ligand (Type 1 or 2) or a stabilizing effector is known (Type 3). Finally, the cutting sites in both GPCRx and GPCRy need to be determined to ensure that the resulting chimera can get activated and successfully transmit the signal from the outside to the inside of the cell. GPCRchimeraDB can come into play to facilitate every stage of this design process, as outlined in the following steps:

#### 1. Explore the entry page of GPCRx

On every entry page, GPCRchimeraDB integrates annotations of major databases in one location, so providing an overview for further analysis. By reviewing known mutations and their impact, the evolutionary score of the protein compared to its subclass, and the conservation of its motifs, a comprehensive picture of crucial residues emerges, so helping to define the sequence space of GPCRx and identifying regions sensitive to mutations. This is critical information, as chimeric design can be seen as introducing stretches of mutations in specific regions. Additional information, such as PTM sites and interacting residues, offer insights on why certain residues might be essential for the protein’s function. The integrated sequence-structure (1D-3D) viewer enables direct examination of these key residues in their 3D context.

#### 2. Define potential GPCRy candidates

The next step is to identify a suitable GPCRy to work with the GPCR under study (GPCRx). For Type 1 and Type 3 chimeras, GPCRy is usually respectively Rhodopsin and the κ-Opioid receptor. However, for Type 2 chimeras, many candidates may be appropriate. The following guidelines can help select the GPCRy:

- Chimeras typically involve combining a well-characterized GPCR with one that is less understood. Therefore, GPCRy should not be an orphan GPCR (this can be established using the ligand-based subclass filter), nor a receptor with limited information on its entry page.
- Since preserving key functional connections in the chimera is crucial, designing chimeras between related GPCRs is in principle more likely to succeed than between distant GPCRs. The filtering system on the GPCRchimeraDB entries list page can be used to identify GPCRs from the same subclass as GPCRx.
- Depending on the research objective, it may be important to select a GPCRy that either binds the same or a different G protein compared to GPCRx. The integrated filtering system can also assist in identifying GPCRs based on their G protein coupling profile.
- GPCRy candidates can also be identified by reviewing past chimeric designs involving GPCRx (see bottom of entry page GPCRx). Related GPCRs of past parents could also be considered.
- GPCRy should be a receptor that is easy to work with in the lab. For instance, opioid related GPCRs may pose practical challenges due to regulatory or experimental constraints.

Once suitable GPCRy candidates have been identified, their entry pages should be explored in the same manner as outlined in Step 1 to gain insights into the key residues for GPCRy’s function.

#### 3. Respect the key rules for determining cutting sites

The cutting sites for GPCRx and GPCRy chimeras need to be carefully determined. While no standardized procedure exists, based on the studied literature we recommend the following guidelines to enhance the success rate of chimera designs:

- Preserve the microswitches essential for the function of the EC parent within the chimera’s sequence ^10,11^.
- Conserve the secondary structure from the parent GPCRs within the chimera by introducing the cutting sites within the parents’ regions with the same secondary structure^11^.
- Favor the EC parent: the more closely the chimeric sequence resembles the EC parent sequence, the better. For example, studies have shown that exchanging IC loop 1 is not critical for successful chimera design^43,51^.
- Ensure that the IC parent contributes its residues responsible for the binding of the G protein of interest to the chimeric sequence^11^.

GPCRchimeraDB supports these rules by providing information on conserved motifs, secondary structures, and interacting residues.

#### 4. Improve the cutting point selection by analyzing related chimeras

Although the above rules are essential for successful chimera design, they may not be sufficient. We suggest reviewing previously designed chimeras involving GPCRx, GPCRy, or related GPCRs, or chimeras designed for the same research purposes. GPCRchimeraDB offers the following assisting features:

- Compare biophysical profiles of previously designed chimeras with their parent GPCRs using the 1D-3D viewer (Figure 2F-G). Does the biophysical profile of the chimera match with the profile of the parents?
- Analyze residue conservation in exchanged regions using the ‘Conservation parent vs chimera’ track or GPCRchimeraDB’s alignment tool. How much do EC parent residues need to be conserved in the provided IC parent region for a chimera to be successful? Is conserving the motifs enough? GPCRchimeraDB also displays the parent subclass conservation: are residues that are conserved throughout the protein subclass required in the chimeric sequence?
- Review the location of the previous cutting sites in the parents and investigate how far they extend into the TM helices.
- Download and align the 3D structures of the parents to identify equivalent regions suitable for positioning the cutting sites.

#### 5. Design the chimera

Once cutting sites have been defined, align GPCRx and GPCRy using the GPCRchimeraDB alignment tool and design your chimera.

To increase the likelihood of success, *in silico* validation is recommended. For example, the AF2 or ESMF model of the designed chimera can be generated and superimposed with the 3D structure of GPCRx and GPCRy to study how much they differ. The model’s confidence metrics (pLDDT and PAE) could also be compared with those of the parent models. Finally, we also suggest computing the biophysical profile of the designed chimera with the b2btools to ensure its properties align with natural GPCR behavior. Molecular Dynamics (MD) simulations could also be considered. The most promising designs can then be tested experimentally before pursuing the research objectives that motivated the chimera’s creation.

## Discussion and perspectives

We present GPCRchimeraDB, a comprehensive database dedicated to both natural and chimeric GPCRs. As the first centralized repository for chimeric GPCRs, it offers extensive, standardized descriptions at the sequence, structural, and biophysical level, complemented by visualization tools and links to their natural, parent GPCRs. This standardization facilitates comparisons between chimeric designs and the parent receptors, providing insights into the requirements for successful designs. To support this, GPCRchimeraDB introduces a naming convention and a classification system based on the type of design (Type 1, 2 or 3). Furthermore, expanding on resources like GPCRdb and UniProt, GPCRchimeraDB enriches the description of natural GPCRs by including their biophysical profile and integrating an interactive sequence-structure visualization tool. By covering all natural GPCRs, it also supports the exploration of novel chimeric designs through new parent combinations. Additionally, the database incorporates a sequence alignment tool, a feature comparator, and a step-by-step workflow to guide users through the chimera design process, making it a valuable resource for both analyzing past designs and developing new ones. Finally, assuming that chimeric GPCRs are functional only if they retain key sequence, structural, and biophysical properties of their parent GPCRs, analyzing previous chimeric designs also brings insights into the key regions of natural parent GPCRs and helps understanding their function.

Given the vast amount of information within GPCRchimeraDB, an AI tool could further streamline the design process. Such a tool could assist researchers by highlighting relevant data and suggesting potential constructs based on their GPCR of interest and research goals. However, to ensure the success of this AI-driven approach, it will be essential to standardize the functionality values of the chimeras or, to provide complete experimental contexts next to each functionality value in order to create a robust dataset for AI training and testing. High-throughput studies under uniform conditions would help to obtain such a dataset.

The strength of GPCRchimeraDB lies in its extensive overview of the GPCR chimera field, providing key insights to guide the design of novel chimeras. We believe this resource will promote a more systematic and informed approach to chimera development, ultimately increasing the success rate of these designs, lowering experimental costs, and deepening our understanding of GPCRs.

## Data availability

The entries and their annotations are directly available on the database (https://www.bio2byte.be/gpcrchimeradb/) or on Zenodo (10.5281/zenodo.14989364). The MA is also available on Zenodo as well as a list of GPCR chimeric designs that did not meet the inclusion criteria for GPCRchimeraDB. The Python scripts to retrieve the annotations to describe the entries as well as to format a chimera’s data based on a sequence are accessible on GitHub (https://github.com/Bio2Byte/GPCRchimeraDB).

## Author contributions

*Charlotte Crauwels:* Data Curation, Formal analysis, Software, Validation, Writing - Original Draft, Funding acquisition. *Adrián Díaz:* Software, Validation. *Wim Vranken:* Conceptualization, Validation, Writing - Review & Editing, Supervision, Funding acquisition.

## Acknowledgments

We thank the Bio2Byte research group for their testing and feedback to help us improve GPCRchimeraDB. We also highly appreciated the feedback provided by Prof. Julien Hanson and his group, as well as by Dr. Sarah Triest. Finally, we would like to express our gratitude to Dr. Zara Sands for her ideas, feedback, and enthusiasm during the early stages of development of this database.

## Declaration of generative AI and AI-assisted technologies in the writing process

During the preparation of this work the authors used ChatGPT (OpenAI) as an editing tool to improve the readability and language of the manuscript. After using this tool, the authors reviewed and edited the content as needed and take full responsibility for the content of the published article.

