## Supplementary material for "GPCRchimeraDB: A database of chimeric G protein-coupled receptors (GPCRs) to assist their design"

**Table S1:** Predicted 3D models available in GPCRchimeraDB.

Between brackets the date of the last update of the template dataset used by the predictor. For ESMF, this date corresponds to the model publication date. AFms is a variant of AF2 without Multiple Sequence Alignment (MSA) input features where activation state-specific GPCR structures are used as templates, so enabling users to generate state-specific models. AFDB stands for AlphaFold Protein Structure Database.

| **Type of GPCR** | **AlphaFold2**^1^ **(AF2)** | **AlphaFold multistate**^2^ **(AFms)** | **ESMFold**^3^ **(ESMF)** |
| --- | --- | --- | --- |
| **Natural** | - If available: From GPCRdb^4^ (2024) - Else: From AFDB^5^ (2022) | Only for human GPCRs:   - If available: From GPCRdb (2024) - Else: From AFms dataset (2023) | Not available in GPCRchimeraDB |
| **Chimeric** | Predicted on VSC Tier-2 general-purpose clusters provided by VUB-HPC | | |
|  | (2024) | (2023) | (2022) |

### Procedure to define the transmembrane limits

The boundaries of the transmembrane (TM) regions were initially assigned based on DSSP^6^ annotations computed from the AF2 models. When available, experimental structures were used instead. However, we observed that AF2 predicts excessively long TMs for certain receptors. For example, TM5 and TM6 of ACM3 are predicted to be 15 to 20 residues longer than in the corresponding experimental structure (PDB ID: 8E9W). To define the TM regions in both chimeric and natural GPCRs, we therefore established minimal and maximal boundaries for each of the 7 TMs (referred to as "min." and "max." TM limits) using the following procedure:

1. We identified 192 natural GPCRs with at least one experimentally resolved 3D structure (as of February 2025). For each GPCR, we selected the highest-resolution structure, irrespective of its activation state and computed its DSSP annotations.
2. The DSSP-derived secondary structure codes were mapped onto their corresponding sequences within a MSA containing these 192 GPCRs (see MA in ‘Alignment procedure GPCRchimeraDB’ for details).
3. For each TM region, we determined the first residue where at least 50% (min. start) and 80% (max. start) of the aligned GPCRs have a DSSP-annotated helix (DSSP codes ‘H,G,I’, Figure S1). Similarly, we defined min. end and max. end as the last positions where at least 80% and 50% of the GPCRs, respectively, retain a helical annotation.

Next, for each GPCR (natural and chimeric), the TM limits were defined as such:

1. DSSP annotations were assigned based on:

- An experimental 3D structure (if available, selecting the highest-resolution structure regardless of activation state).
- Otherwise, AF2 models were used (either from GPCRdb, AFDB or predicted in-house, see section ‘Structural description’ in the main text).

2. If a continuous stretch of at least 10 residues was predicted as helical, it was classified as a TM region. These helices were then refined if needed to obtain 7 TM regions in total:

- If a helical region starts before min. start, it is trimmed to begin at min. start. If it starts after max. start, it is extended to begin at max. start. If it starts between min. and max. start, it is not modified.
- The end of the TMs is adjusted similarly using the min. and max. end values.
- If an entry has two helical regions within the min. start and max. end of a TM (e.g., due to a kink), they are merged into a single TM region.

As shown in Figure S2, this results in TM regions for the 1755 natural entries that align well with those observed in the 192 unique experimental structures, even allowing for multiple helical turn variations between the receptors. This validates the robustness of our approach on natural GPCRs and allows us to use it to define the TM limits of the chimeric GPCRs in GPCRchimeraDB.

For chimeric entries, the procedure remains the same, except for the definition of the min. and max. TM limits, as chimeric sequences are not part of the MSA used to establish these boundaries.
To assign TM limits to chimeric sequences, we first align each chimeric sequence to its EC parent, which is included in the MSA. This alignment allows us to transfer the min. and max. TM limits from the EC parent onto the chimeric sequence, ensuring consistency with the TM boundaries defined for natural GPCRs.

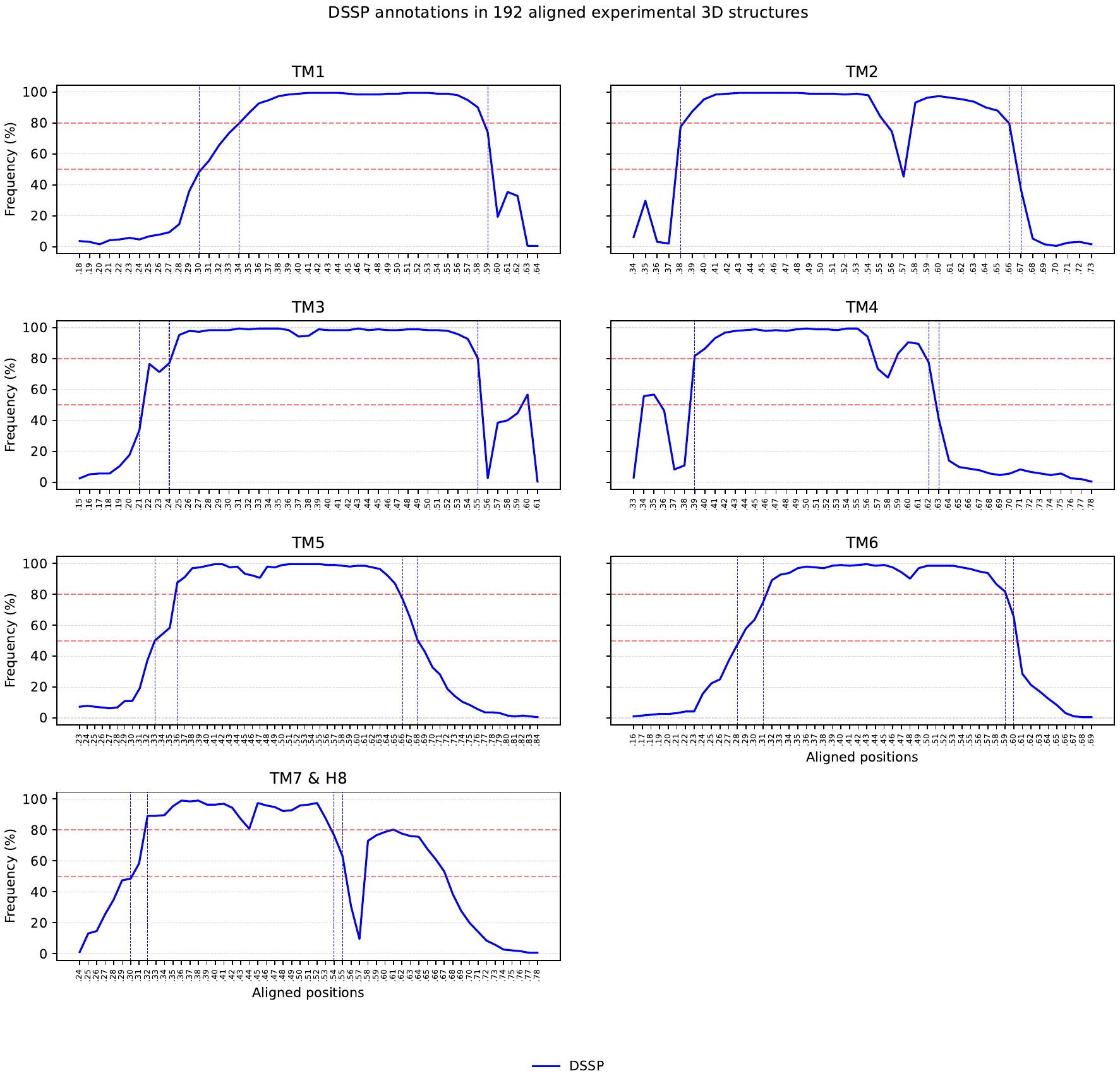

**Figure S1**: Frequency of residues predicted helical in aligned natural GPCRs.

DSSP annotations were computed for 192 unique experimental 3D structures of natural mammalian class A GPCRs and mapped on their aligned sequences. For each TM region, we identified the first residue where at least 50% (min. start) and 80% (max. start) of the aligned GPCRs exhibit a DSSP-annotated helix (code ‘H,G,I’). Similarly, the min. end and max. end correspond to the last positions where at least 80% and 50% of GPCRs, respectively, retain a helical annotation These limits are used to refine the TM limits of the GPCRs in GPCRchimeraDB if their helical regions are predicted to be longer or shorter than these limits.

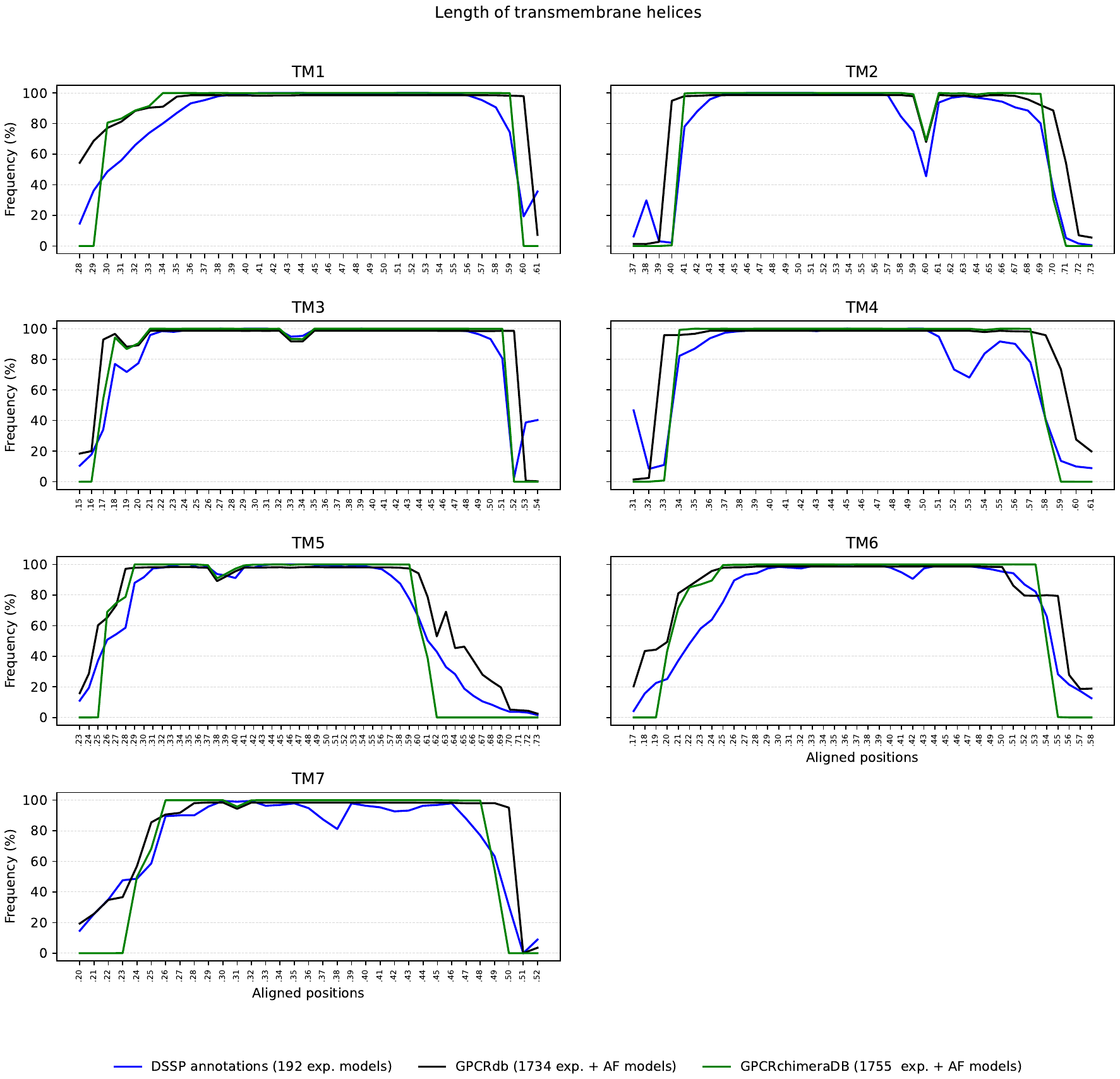

**Figure S2:** Length distribution of the TM helices.

Depending on the methodology used to determine the TM helix limits, the length can vary. GPCRdb’s (black) and GPCRchimeraDB’s (green) methodology align well with the experimental data (blue).

### Alignment procedure GPCRchimeraDB

To generate an alignment	(<https://www.bio2byte.be/gpcrchimeradb/sequence_alignment_entries>), the sequence alignment tool in GPCRchimeraDB operates in the following steps:

1. **Selecting sequences to align:** On the sequence alignment page of GPCRchimeraDB, users select proteins to align from the list of entries of the database (referred to as ‘mandatory sequences’). Additionally, users may input or upload their own sequences in FASTA format (referred to as ‘optional sequences’). A minimum of two mandatory sequences must be selected.
2. **Chimeric protein handling:** If chimeric proteins are included among the mandatory sequences, the parent sequences of these chimeras are automatically identified and added to the mandatory sequences dataset for alignment.
3. **Retrieval of the pre-aligned natural sequences:** For the natural proteins in the mandatory sequence dataset, the tool retrieves their aligned sequences from a pre-computed Master Alignment (MA, see below).
4. *If the dataset includes chimeric proteins and/or optional sequences:*

- **Aligning chimeric or optional sequences:** A Hidden Markov Model profile (pHMM) is generated based on the aligned sequences from step 3 using HMMER^7^ (via the ’hmmbuild’ command). This pHMM is then used to align these chimeric and/or optional sequences (via the ’hmmalign’ command).

*If the dataset does not include chimeric proteins or optional sequences:*

- **The subset of the MA is the final MSA:** The final alignment will consist of the extracted already aligned sequences from step 3.

5. **Final MSA Visualization and Download:** The resulting MSA, in FASTA format, includes natural sequences and the eventual chimeric sequences and their parents and/or the optional sequences. Users can visualize and download the MSA, which is colored according to the CLUSTAL scheme. Secondary structure annotations are based on the consensus aligned secondary structure annotation of the proteins in the MSA.

The MA used in this procedure was constructed as follows:

1. **Structure-guided alignment of all human GPCRs:** A MSA of all natural human class A GPCRs (711 sequences, InterPro IPR017452) was generated using the structure-informed MSA pipeline SIMSApiper^8^. Structural templates guiding the alignment were selected based on the following criteria:
   - If an experimental 3D structure was available for a given GPCR, it was retrieved. When multiple structures existed, the one with the highest resolution was selected, regardless of activation state. A total of 181 structures were obtained.
   - If no experimental structure was available, the AlphaFold2 model of the GPCR was retrieved:
     - From GPCRdb, if available (updated templates from 2024, 85 models downloaded).
     - Otherwise, from AFDB (templates updated in 2022, 445 models downloaded).

The specific SIMSApiper parameters applied are detailed below.

2. **Incorporation of non-human mammalian GPCRs:** Natural non-human mammalian class A GPCRs (1,044 sequences) were added to the initial MSA using the MAFFT ‘add’ function, with G-INS-1 as the alignment strategy. The alignment was further refined by squeezing the loop regions relative to the conserved TM helices. This refinement is based on the principle that the spatial position of amino acids in relation to conserved secondary structure elements is more significant than their exact sequence identity^9^. The positions of the helices were determined using DSSP annotations from the 3D structures. For 11 non-human mammalian class A GPCRs their experimental 3D structure was used as structural information. For the remaining GPCRs, their AlphaFold2 model from AFDB was used.

The final MA is available on Zenodo (10.5281/zenodo.14989364) together with the used 3D structures and models.

The SIMSApiper parameters used to compute the MA were the following:

nextflow run simsapiper.nf \

-profile supercomputer \

--data $data/data \

--createSubsets 30 \

--minSubsetID “min” \

--seqQC 5 \

--retrieve true \

--strucQC 1 \

--dssp true \

--squeeze ‘H’ \

--squeezePerc 70 \

--reorder true \

--outFolder $output_folder \

### Alignment GPCRchimeraDB vs GPCRdb: minor but critical differences

As described by Kooistra et al.^10^, GPCRs lacking an experimental 3D structure are aligned in GPCRdb by inferring the start and end of secondary structure elements from receptors with similar motifs, lengths or overall sequence, i.e. they are aligned purely based on their sequence information. Nevertheless, with the development of AlphaFold2, structural information is now available for all proteins. As shown by Baltzis et al.^11^, even predicted models of reduced accuracy can significantly improve sequence alignments. Motivated by this, we generated a new structure-informed MSA of all human GPCRs (extended to all mammalian GPCRs) and compared it to the MSA provided by GPCRdb.

While both alignments are in agreement for most receptors (see <https://www.bio2byte.be/gpcrchimeradb/sequence_alignment_entries>), notable differences emerge for some orphan GPCRs. These discrepancies can be attributed to the lack of structural data and the low sequence identity of these receptors compared to well-characterized families. For example, TM5 in the SREB GPCRs (GPR27/85/173) as well as in GP148 and GP149 is misaligned in GPCRdb (Figure S3A-B), and the same is true for TM2 in GPR146 (Figure S3C). These receptors share less than 15% sequence identity with Rhodopsin.

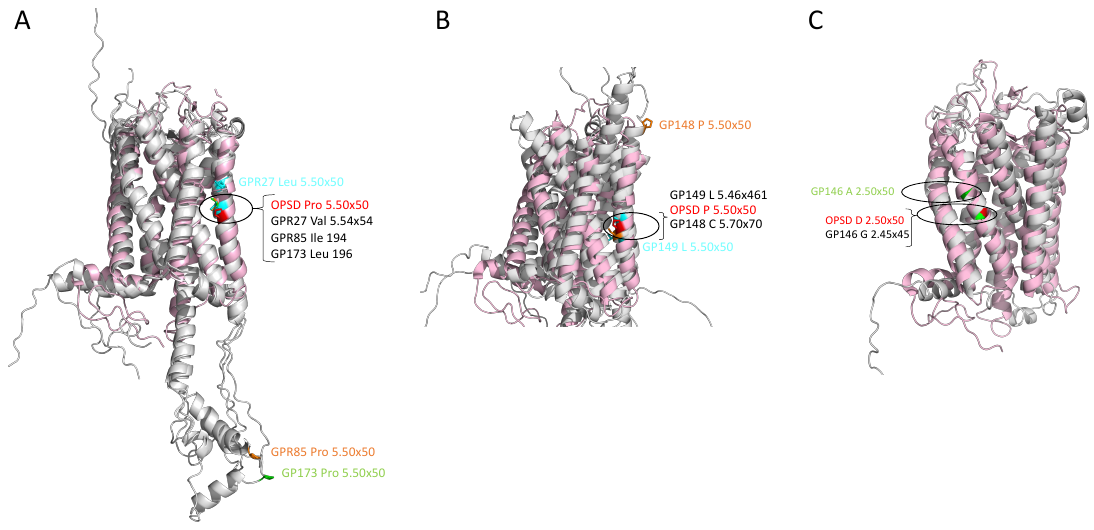

**Figure S3:** Structural alignment of orphan GPCRs with Rhodopsin (PDB: 5TE5, shown in pink).

The residues aligning with the x.50 position of Rhodopsin (in red) are indicated. Notably, these residues do not correspond to the annotated x.50 positions of the orphan GPCRs (in green, orange and/or cyan). Since the numbering is based on GPCRdb annotations, this discrepancy reveals a misalignment in the GPCRdb-derived sequence alignments. In contrast, GPCRchimeraDB's alignment is consistent with the structural alignment shown here. **A.** GPR27, GPR85, and GP173 (in grey) aligned to Rhodopsin. **B.** GP148 and GP149 (in grey) aligned to Rhodopsin. **C.** GPR146 (in grey) aligned to Rhodopsin.

Because our alignment and TM region definition can slightly differ from those used in GPCRdb, some discrepancies in the residue numbering may appear for some positions, particularly in orphan receptors. For example, a residue annotated as part of a loop region in GPCRchimeraDB (e.g. P233 GPR85 located in ICL3) may correspond to a helical residue in GPCRdb (P5.50x50, see Figure S3). To maintain compatibility with established resources, we have chosen to retain the original BW and GPCRdb annotations within GPCRchimeraDB, without modifying them to fit our own alignment. Where such inconsistencies occur, we clearly flag them in the database interface, including on entry-specific pages and the alignment results page. This allows users to interpret these differences and make informed choices. In the long term, we support community-driven efforts to harmonize the annotation of orphan GPCRs and such updates can then be integrated into future versions of GPCRchimeraDB.

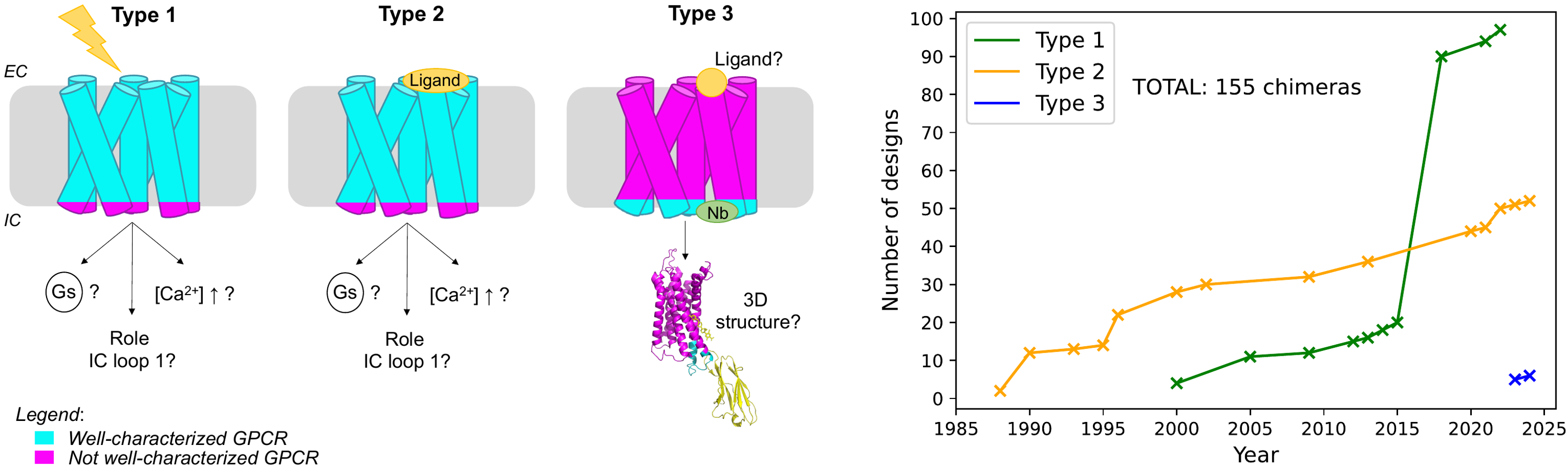

**Figure S4:** Schematic illustration of chimeric design types in GPCRchimeraDB.

Type 1 chimeras, also known as OptoXRs, are activated by light stimulation. Type 2 chimeras are similar in design to Type 1 but are activated by endogenous or synthetic ligands instead of light. Type 3 chimeras incorporate a stabilizing effector, such as a nanobody or VHH, that locks the GPCR in a specific conformation. The choice of chimera design depends on the research objective. Type 1 and Type 2 chimeras are typically used to investigate GPCR signaling pathways or to identify key regions critical for protein function. In contrast, Type 3 chimeras are designed for identifying potential ligand binders or supporting 3D structure determination.

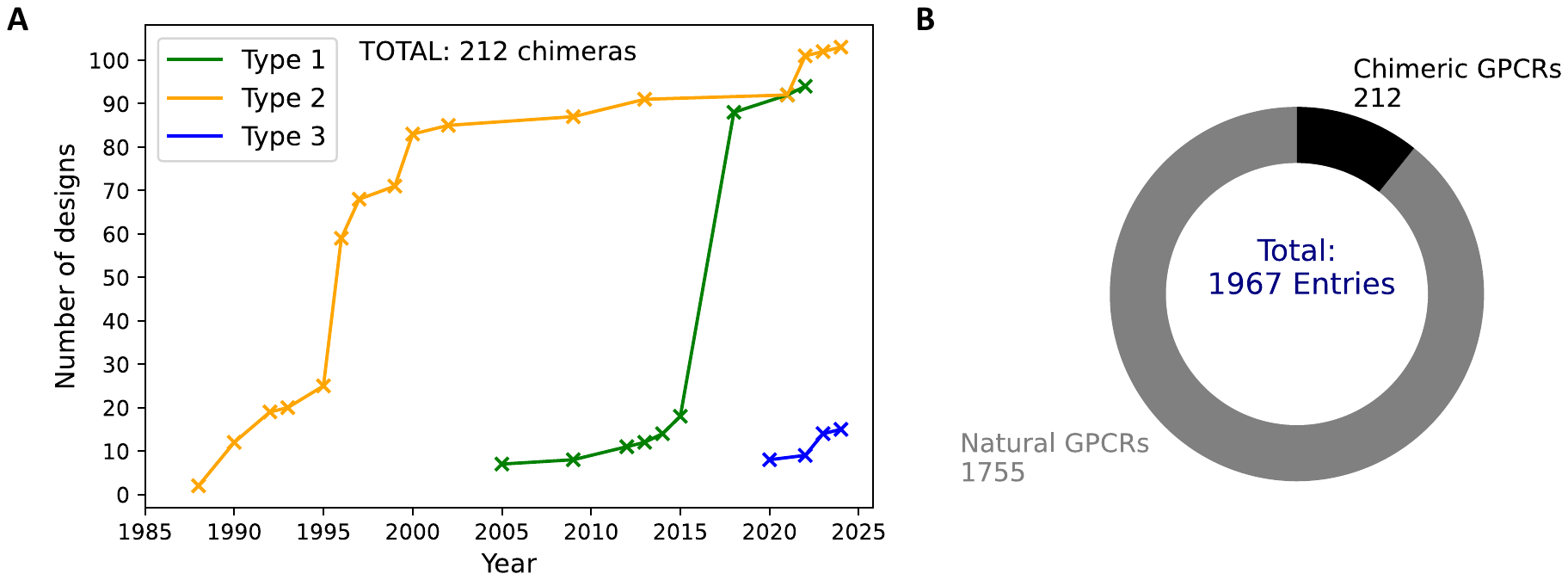

**Figure S5:** Data in GPCRchimeraDB.

**A.** Cumulative number of GPCR chimeras by year of publication as available in GPCRchimeraDB. **B.** Distribution of natural and chimeric GPCRs currently available in GPCRchimeraDB.

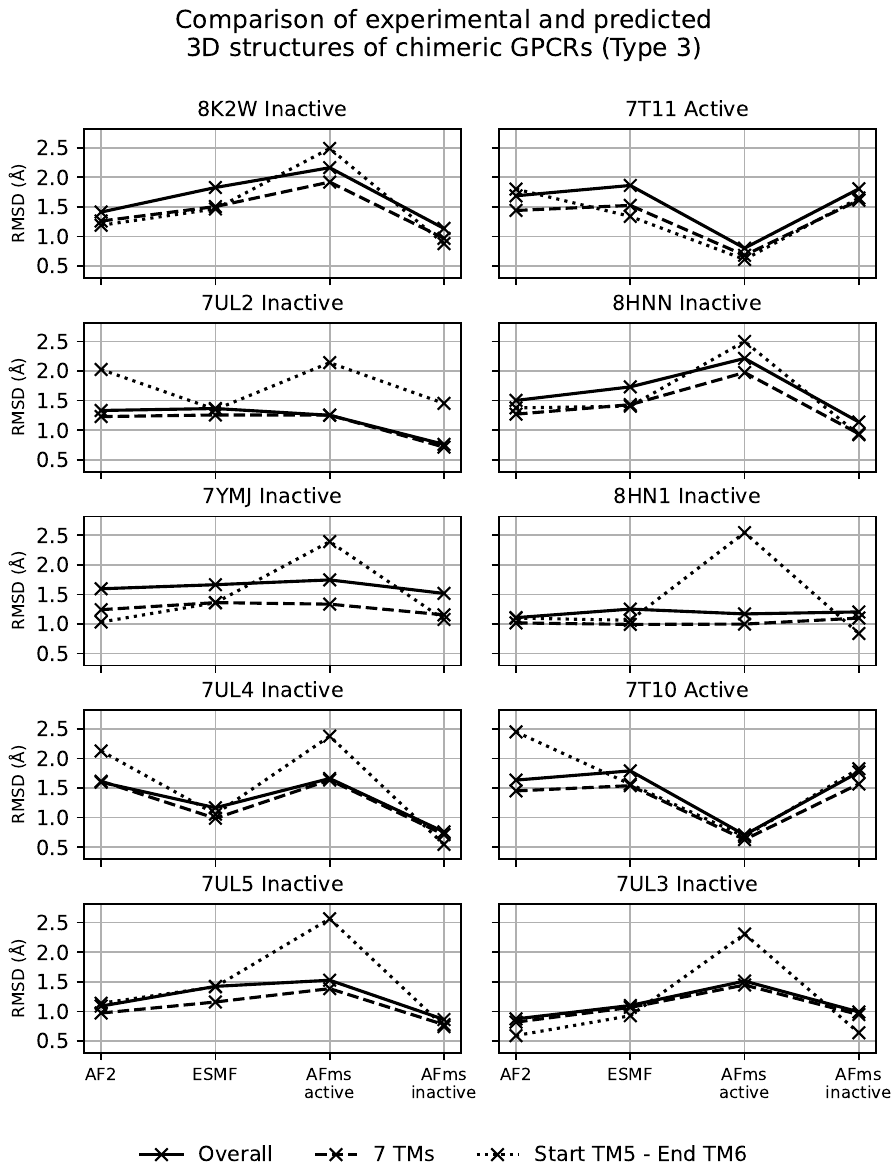

**Figure S6:** Comparison of the experimental and predicted 3D structures of Type 3 chimeras.

The 10 PDB^12^ entries (from 7 unique chimeras) used in this analysis are the only experimentally resolved chimeric GPCR structures available as of February 2025. Structural alignment was performed using PyMOL’s ‘align’command. All chimeras share 2 common features: a single swapped region located around the end of TM5 and the beginning of TM6 and, their IC parent is κ opioid receptor.

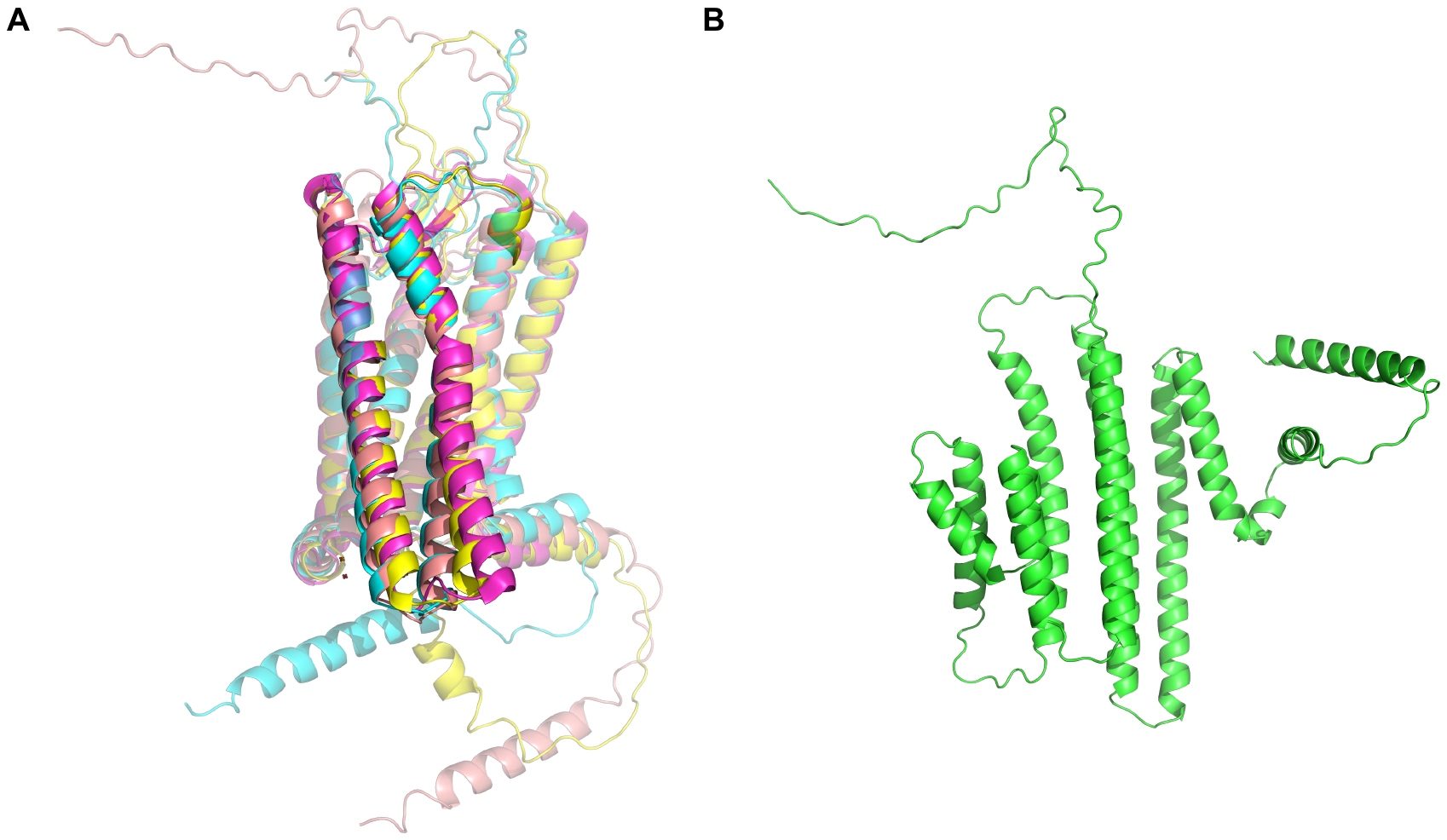

**Figure S7:** Experimental 3D structure and predicted models of a chimera.

The chimera analyzed, SSR2_HUMAN_OPRK_HUMAN_1, was constructed by replacing the intracellular loop 3 (ICL3) and beginning of TM6 of the Somatostatin type 2 receptor with the analogous regions of the κ-opioid receptor. **A.** In magenta the experimental 3D structure (7T10, active state, PDB), in cyan the AF2 model, in yellow the AFms active state model and in salmon, the ESMF model. Here the AFms active state model is the most accurate (RMSD of 0.669 Å with PDB structure) compared to the AF2 model (1.535 Å) and the ESMF model (1.740 Å). A longer TM5 and misoriented TM6 are common features of predicted models compared to their experimental counterparts. **B.** The predicted model obtained with AF2 single sequence mode^13^. The model is notably inaccurate (see relative position helices compared to **A.**).
